# Label-Free Live Cell Type Prediction by Integrating Raman Spectroscopy and Machine Learning

**DOI:** 10.64898/2026.06.16.732770

**Authors:** Adrian Lita, Noor E. Zannat, Helena Muley, Nicoleta Siminea, Sophia Spinu, Joel Sjoberg, Andrei Paun, Yuri Nikulin, Christel Herold-Mende, Ion Petre, Mioara Larion

**Affiliations:** Neuro-Oncology Branch, Center for Cancer Research (CCR), National Cancer Institute, National Institutes of Health, Bethesda, Maryland, USA; Department of Mathematics and Statistics, University of Turku, Finland; Faculty of Mathematics and Computer Science, University of Bucharest, Romania; Division of Experimental Neurosurgery, Department of Neurosurgery, Medical Faculty Heidelberg, Heidelberg University, 69120, Heidelberg, Germany

**Keywords:** machine learning, spontaneous Raman spectroscopy, glioma, cell types, cell fingerprinting

## Abstract

Coherent Raman spectroscopy enables label-free biochemical fingerprinting of live cells with subcellular resolution. We previously developed a machine learning framework capable of classifying glioma FFPE tissues using Raman spectral signatures. To accelerate live cell acquisition, we previously developed RADAR (Raman Spectral Analysis Using Deep Learning for Artifact Removal), a method that increases imaging speed by an order of magnitude while preserving spectral integrity. By integrating high-speed Raman imaging with supervised machine learning, we aimed to define unique biochemical fingerprints specific to cell type. We hypothesized that intrinsic biochemical composition alone is sufficient to distinguish cellular identity and tumor subtype. To test this, we generated metabolic maps of diverse brain-derived cell types—including astrocytoma, oligodendroglioma, and glioblastoma cells—using coherent Raman spectroscopy at single-cell resolution. Patient-derived brain tumor cell lines representing genetically heterogeneous backgrounds were analyzed. Samples were stratified by IDH1 mutation status (IDH1-mutant and IDH1–wild-type) and histologically classified as oligodendroglioma or astrocytoma. Raman spectral data were acquired from 286 live single cells across the two principal molecular classes, with further subdivision into two histologic subtypes within the IDH1-mutant group. Classification was performed using an XGBoost model with shallow tree depth (1–3), a 20% held-out test set, and grouped, stratified 5-fold cross-validation to control for sample-level bias. The machine learning framework distinguished IDH1-mutant from IDH1–wild-type cells with a ROC–AUC of 0.78 and further discriminated IDH1-mutant astrocytoma from oligodendroglioma cells with a ROC–AUC of 0.81. Feature importance analysis demonstrated that separation between IDH1-mutant and IDH1–wild-type cells was driven primarily by Raman peaks associated with protein amide bands, total NADH, unsaturated fatty acids, and heme-related vibrational modes. Within the IDH1-mutant class, discrimination between oligodendroglioma and astrocytoma was driven by lipid-rich vesicle signatures, protein/polyamide amide bands, and lipid-associated spectral features. Together, these findings support the feasibility of label-free, machine learning–assisted Raman profiling to resolve clinically relevant glioma subtypes at single-cell resolution. This scalable analytical framework provides a translational platform for investigating metabolic heterogeneity, therapeutic response, co-culture systems, and patient-derived organoid models.

## Introduction

Accurate molecular classification of cancer cells is essential for understanding tumor biology, predicting therapeutic response, and guiding precision medicine approaches^1,2^. Current methods for molecular characterization typically rely on genomic, transcriptomic, or immunohistochemical analyses that require cell fixation, tissue processing, or destructive sample preparation^2–4^.

In the context of glioma, contemporary molecular diagnostic approaches, including DNA methylation profiling and next-generation sequencing (NGS), have transformed the classification of diffuse gliomas by enabling comprehensive genomic and epigenetic characterization^5–12^. These advances have refined diagnostic accuracy, improved prognostic stratification, and facilitated the development of more personalized treatment strategies. However, despite increasingly sophisticated molecular classification systems and the widespread use of multimodal therapy consisting of maximal safe surgical resection followed by radiotherapy and temozolomide chemotherapy, disease recurrence remains universal^13–15^. This highlights a critical limitation of current approaches: while genomic and epigenetic profiling provide valuable information regarding tumor identity, they do not necessarily capture the dynamic functional states that ultimately govern cellular behavior, therapeutic response, and disease progression, because they provide only static snapshots of cellular states and are not readily adaptable to longitudinal monitoring of living cells. Emerging evidence suggests that metabolic reprogramming, cellular plasticity, and lineage-specific biological programs contribute substantially to glioma heterogeneity and treatment resistance^16–19^. Consequently, there is growing interest in technologies capable of interrogating the biochemical consequences of genetic alterations directly in living cells.

Coherent Raman spectroscopy offers a unique opportunity to address this challenge. As a label-free and non-destructive technique, Raman spectroscopy measures intrinsic molecular vibrations arising from proteins, lipids, nucleic acids, carbohydrates, and metabolites, thereby providing a biochemical fingerprint of cellular phenotype. Because spontaneous Raman spectra arise from intrinsic molecular vibrations, they provide a biochemical fingerprint reflecting the abundance and organization of proteins, lipids, nucleic acids, carbohydrates, and metabolites within individual cells^20–26^. Unlike fluorescence-based approaches, Raman spectroscopy does not require exogenous labels or genetic modification, preserving cellular integrity and enabling repeated measurements of living cells over time. These properties make Raman spectroscopy particularly attractive for studying dynamic biological processes and cellular heterogeneity.

Recent advances in instrumentation, computational analysis, and machine learning have significantly expanded the utility of Raman spectroscopy in biomedical applications^27–29^. Raman-based approaches have been successfully used to distinguish normal and malignant cells, classify tumor subtypes, identify stem-like populations, monitor drug responses, and characterize metabolic states in cancer types^24,25,30–39^. Importantly, because Raman spectra capture integrated biochemical information rather than individual molecular markers, they can reveal functional cellular phenotypes that may not be apparent from genomic alterations alone.

Cancer-associated mutations such as R132H mutation in IDH1 in glioma, frequently induce profound metabolic reprogramming that alters cellular composition and organelle function^6,9,40–44^. These biochemical changes can influence the abundance of lipids, proteins, metabolites, and redox-active molecules, generating spectroscopic signatures that are detectable by Raman analysis. In particular, alterations in mitochondrial metabolism, lipid synthesis, and cellular redox state are increasingly recognized as important determinants of tumor behavior and therapeutic sensitivity^45^. As a result, Raman spectroscopy offers a unique opportunity to interrogate these functional consequences directly in living cells. In the context of diffuse gliomas, such an approach may be particularly valuable because key molecular alterations, including IDH1 mutations, exert profound effects on cellular metabolism, mitochondrial function, redox homeostasis, and lipid organization^9,22,40,41,44,46–54^. These downstream biochemical consequences are potentially detectable by Raman spectroscopy and may provide complementary information beyond genotype alone. Therefore, live-cell Raman spectroscopy represents a promising platform for linking molecular alterations to their functional phenotypic outcomes and for identifying biologically meaningful states that may influence therapeutic vulnerability and clinical behavior.

In this study, we investigated whether spontaneous Raman spectroscopic profiling of living cells could accurately predict molecular status based solely on intrinsic biochemical signatures. By integrating live-cell spontaneous Raman measurements with computational classification approaches, we sought to identify the spectral features underlying prediction and to determine the biological processes that distinguish the analyzed cellular populations. Our findings demonstrate that label-free Raman spectroscopy captures reproducible biochemical signatures associated with molecular phenotype and highlight its potential as a non-destructive platform for real-time cellular classification.

## Materials and Methods

### Cell lines and Raman data

GSC923 (GBM) were developed by the Neuro-Oncology Branch (National Cancer Institute). TS603 (oligodendroglioma) were kindly provided by Dr. Timothy Chan’s lab; NCH612 (oligodendroglioma), NCH551B (astrocytoma) and NCH1681 (astrocytoma) by Dr. Christel Herold-Mende’s lab; and L0 (GBM) and L1 (GBM) by Dr. Jinkyu Jung. BT237 (oligodendroglioma) was received from Oligo Nation. BT142 (oligoastrocytoma), U87 MG cells (GBM) and 293T were purchased from ATCC and U251 MG (GBM) from Sigma. U251 MG cells, U87 MG and 293T cells were cultured in DMEM (10-013-CV Corning) supplemented with 10% fetal bovine serum (SH12450H R&D Systems) and 1% penicillin-streptomycin (15140122 Gibco).

All the experiments were performed using human-derived attached cells. Neurospheres were grown in suspension in DMEM/F12 medium (11320033 Gibco) supplemented with 1% N2 growth supplement (17502048 Gibco), 2 μg/mL heparin sodium salt (07980 Stem Cell), 20 ng/mL EGF (236-EG R&D Systems), 20 ng/mL FGF (3718-FB R&D Systems) and 1% penicillin-streptomycin (15140122 Gibco). Two days before the experiment the cells were plated on a 35 mm diameter dish (Ibidi GmbH) on media containing 10% FBS (R&D Systems) which allowed the cells to attach to the plate. The day of the measurement, the media was changed to replace DMEM/F12 to FluoreBrite (A1896701 Thermofisher Scientific).

The dataset consisted of live-cell Raman measurements acquired from several glioma-related cell lines and normal cells. The analyzed samples included 29 oligodendroglioma samples, 18 astrocytoma samples, 208 glioblastoma (GBM) samples with wildtype IDH1 status, 20 GBM samples carrying a synthetically engineered IDH1 mutation, and 6 normal-cell samples. In total, the dataset contained 698,193 Raman spectra.

Raman measurements were acquired as two-dimensional maps covering individual cells or small groups of cells. Most maps consisted of 50 × 50 measurement locations, resulting in a similar number of spectra per sample. The number of spectra therefore greatly exceeded the number of biological samples. Since spectra originating from the same sample are not independent observations, all model validation procedures were performed at the sample level.

### Spectral preprocessing

Raw Raman spectra were preprocessed using RADAR (Raman Spectral Analysis Using Deep Learning for Artifact Removal), a deep-learning framework designed for simultaneous correction and denoising of Raman spectra. RADAR decomposes each spectrum into four components corresponding to the baseline, cosmic rays, noise, and Raman peaks^27,28^. The models were trained using synthetically generated Raman spectra with known ground-truth components, allowing supervised learning of spectral artifact removal.

For each acquired spectrum, RADAR estimated the baseline, cosmic-ray, noise, and peak contributions. The reconstructed peak component was retained for all subsequent analyses. This procedure removes fluorescence-related baseline distortions, sporadic cosmic-ray artifacts, and measurement noise in a single processing step while preserving the spectral features associated with the biochemical composition of the sample.

The use of RADAR also ensured a consistent preprocessing procedure across all samples and classification tasks. The resulting peak spectra were subsequently used for cellular-region identification and machine-learning analysis.

### Identification of cellular regions

Each Raman map contained spectra originating both from cellular material and from the surrounding medium. Since spectra from the medium contain limited information about cellular phenotype, an additional filtering step was performed before classification.

For each Raman map, k-means clustering was applied to partition spectra into groups with similar spectral characteristics. The resulting clusters were inspected and assigned either to cellular regions or to background regions dominated by water and medium signals. Only spectra belonging to cellular regions were retained for further analysis.

### Classification tasks

Several binary classification tasks were considered to evaluate the ability of Raman spectra to distinguish between different cellular phenotypes and molecular states. The analyzed tasks were:

- Astrocytoma versus oligodendroglioma;
- GBM wildtype (GBM^WT^) versus oligodendroglioma;
- GBM wildtype (GBM^WT^) versus astrocytoma and oligodendroglioma combined (patient IDH1^mut^);
- GBM IDH1 mutant (GBM^mut^) versus GBM IDH1 wildtype (GBM^WT^);
- Normal cells versus oligodendroglioma;
- Normal cells versus astrocytoma.

The comparisons involving normal cells were included as exploratory analyses and were not considered the primary focus of the study.

### Machine-learning model

Classification was performed using the XGBoost framework. For all experiments, the model consisted of 400 decision trees with a maximum depth of 2 and a binary logistic objective function. To account for substantial differences in class sizes, class weights were incorporated during training. The same model configuration was used for all classification tasks.

### Validation procedure

Model performance was evaluated using grouped five-fold cross-validation. All spectra originating from the same biological sample were assigned to the same fold, ensuring complete separation between training and test samples.

The classifier generated predictions at the spectrum level. For each sample, a final class assignment was obtained using majority voting across all spectra belonging to that sample. Performance metrics were then computed using these sample-level predictions.

Classification performance was assessed using balanced accuracy, precision, recall, F1-score, and the area under the receiver operating characteristic curve (ROC AUC).

Another validation was performed in model co-culture of two cell lines; one cell lines was labeled with DAPI while the other was cultured unlabeled. The cells were allowed to attach, and Raman spectra of the two cell lines was recorded. Then, the data was submitted to machine learning for prediction.

## Results

### Workflow for Label-Free Prediction of Glioma Cell Identity from Live-Cell Raman Spectra

The molecular signatures associated with a cellular identity, metabolic or proliferative status, are encoded within their spontaneous Raman spectra and can be decoded by machine learning models to enable label-free prediction of cell identity and cellular state. To determine whether intrinsic biochemical signatures captured by Raman spectroscopy can be used to predict glioma cell identity, we analyzed a panel of patient-derived glioma cell lines initially established from surgically resected tumors (Fig. 1). Cells were initially maintained as neurospheres and subsequently transitioned to adherent culture conditions to facilitate live-cell Raman imaging.

**Figure 1.**
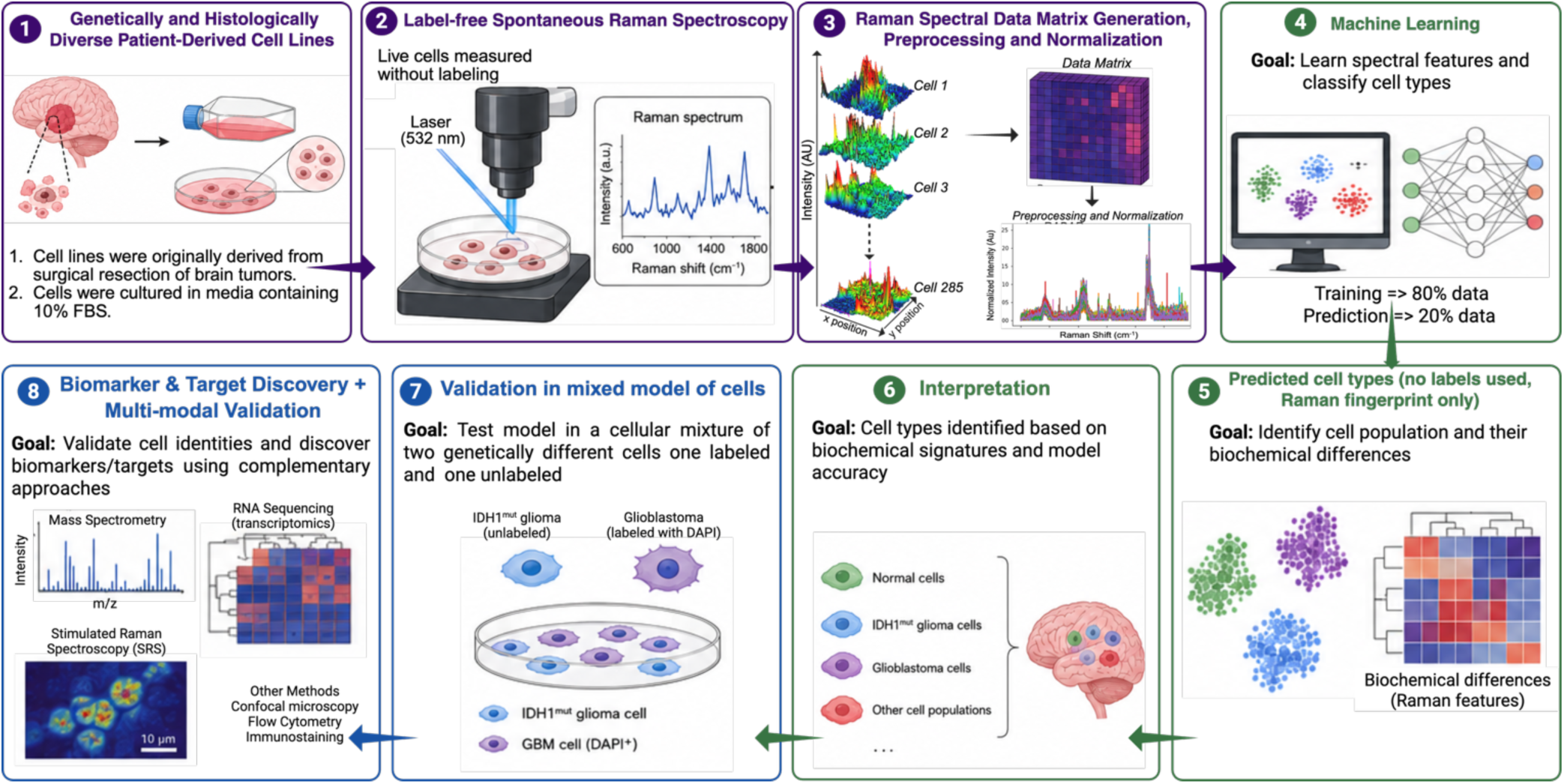
The overview of the study design. 1. Cell lines that were derived from different surgically resected tumors were cultured in media as neurospheres. After enough cells were present, media was changed to allow the cells to attach. **2.** Live cells were measured using spontaneous Raman spectroscopy. **3.** After cells were acquired, the data was exported for pre-processing using our developed RADAR method, normalized and exported for machine learning. **4.** Data was split into 80 % training data set and 20% prediction. Classification models were trained using grouped five-fold cross-validation and evaluated at the sample level using majority voting across all spectra belonging to a sample. **5.** Predicted cell types were compared with their actual class and Raman spectral features were extracted to identify biochemical differences. **6.** Different two class comparison was conducted to identify biochemical differences. **7.** Validation was conducted using mixed population from each comparison (IDH1^mut^ versus IDH1^WT^) in which one cell line was labeled with DAPI for class identification in the mixture. This mixture was subjected to the classification. **8.** Biomarkers associated with one cell type were identified as well as biochemical targets for future validation and multi-modal analyses.

Live-cell Raman hyperspectral maps were acquired over 50 µm × 50 µm regions, encompassing either single cells or small multicellular clusters. The resulting three-dimensional Raman datasets contained spectral contributions from both the culture medium and intracellular components. Leveraging the spatially resolved nature of the measurements, medium-derived spectra were distinguished from cell-associated spectra and removed prior to analysis (Fig 1. Panel 3). This preprocessing step enabled subsequent machine learning models to focus exclusively on intracellular biochemical fingerprints relevant to cell-type classification. The remaining cellular spectra were normalized and used for machine learning–based classification. To isolate intracellular biochemical signals, Raman datasets were processed using our RADAR pipeline^27,28^. To evaluate whether Raman-derived biochemical fingerprints were sufficient to distinguish genetically distinct glioma populations, the dataset was partitioned into training (80%) and independent test (20%) cohorts. Classification models were trained using grouped five-fold cross-validation and evaluated at the sample level through majority voting across all spectra associated with a given sample. The resulting models accurately predicted cell identity based solely on label-free Raman signatures, demonstrating that intrinsic biochemical composition contains sufficient information to discriminate among glioma cell types (Fig 1. Panel 4).

In the next step, to investigate the molecular basis of classification, Raman features contributing to model performance were examined. Distinct spectral signatures were observed between biologically relevant groups, including IDH1^mut^ and IDH1^WT^ glioma cells, revealing biochemical differences associated with specific genetic backgrounds (Fig 1. Panel 5-6). Comparative analyses identified multiple spectral regions that contributed to class separation, suggesting alterations in cellular metabolism, protein composition, lipid content, and nucleic acid abundance.

To assess model robustness under conditions that more closely resemble cellular heterogeneity encountered in vivo, validation experiments were performed using mixed-cell populations generated from pairwise comparisons. In these experiments, one cell population was labeled with DAPI to provide independent ground-truth identification within the mixture. Raman spectra acquired from heterogeneous cultures were subjected to classification using models trained on pure populations. Accurate identification of individual cell types within mixed populations demonstrated that Raman-derived biochemical fingerprints remain sufficiently distinct for cell-type prediction in complex cellular environments (Fig 1. Panel 7).

Finally, discriminative Raman features identified by the classification framework were leveraged to nominate candidate biomarkers associated with specific glioma cell states. These spectral signatures also highlighted putative biochemical pathways and molecular targets that warrant further investigation using orthogonal validation approaches, including mass spectrometry–based proteomics and metabolomics, transcriptomic profiling, stimulated Raman imaging, and other multimodal analytical platforms (Fig 1. Panel 8).

### Raman-based discrimination of glioma cell populations

We first evaluated whether Raman spectra acquired from live-cell measurements could distinguish between different glioma-related cellular populations. Classification models were trained using grouped five-fold cross-validation and evaluated at the sample level using majority voting across all spectra belonging to a sample.

Table 1 summarizes the classification performance obtained for all investigated tasks. Because of the substantial class imbalance present in several comparisons, balanced accuracy was used as the primary performance measure. ROC AUC and PR AUC are reported as complementary metrics.

**Table 1.** Classification performance for the analyzed cell line comparisons.

| Classification task | Balanced Accuracy | ROC AUC | PR AUC |
| --- | --- | --- | --- |
| Astro vs Oligo | 0.73 | 0.78 | 0.80 |
| GBM<sup>WT</sup> vs Oligo | 0.77 | 0.84 | 0.44 |
| GBM<sup>WT</sup> vs Astro + Oligo | 0.62 | 0.68 | 0.33 |
| GBM<sup>mut</sup> vs GBM<sup>WT</sup> | 0.81 | 0.86 | 0.98 |
| IDH1<sup>WT</sup> vs IDH1<sup>Mut</sup> | 0.70 | 0.77 | 0.48 |
| Normal vs Oligo* | 0.75 | 0.87 | 0.96 |
| Normal vs Astro* | 0.81 | 0.92 | 0.82 |

\* Exploratory comparisons due to the limited number of normal-cell samples.

Across the primary classification tasks, balanced accuracies ranged from 0.62 to 0.81. The highest performance was obtained for the comparison between GBM cells carrying an engineered IDH1 mutation and GBM wildtype cells. Comparisons involving oligodendroglioma and astrocytoma cells also showed consistent separation, indicating that Raman spectra contain information related to cellular phenotype and tumor lineage.

The most challenging task was the discrimination between GBM wildtype cells and the combined astrocytoma–oligodendroglioma group. Although performance was lower in this setting, the results remained above random expectation and suggest the presence of measurable biochemical differences between the cellular populations.

### Raman Spectroscopy Reveals IDH1 Mutation–Driven Biochemical Reprogramming

One of the main objectives of the study was to determine whether Raman spectroscopy can detect biochemical changes associated with IDH1 status. To address this question, classifiers were trained to distinguish cells carrying an engineered IDH1 mutation from their wildtype counterparts. The comparison between these two isogenic cell lines namely, GBM mutant (GBM^mut^) and GBM wildtype (GBM^WT^), achieved a balanced accuracy of 0.81, with ROC AUC and PR AUC values of 0.86 and 0.98, respectively. These results indicate that the engineered mutation is associated with reproducible changes in the Raman spectra of live GBM cells.

Feature importance analysis identified Raman bands centered at approximately 3179, 3023–3139, 1627–1633, 1542–1544, 1120–1139, and 867–886 cm⁻¹ as major contributors to the classification of GBM^mut^ and GBM^WT^ cells (Table 2).

**Table 2.** Results from GBM^WT^ versus GBM^mut^ classification.

| Raman Frequency | Normalized Feature Importance | Biochemical Assignments | Molecular Vibrations |
| --- | --- | --- | --- |
| 1627.50529 | 1 | Proteins (Amide I) | C=O stretching |
| 1120.31852 | 0.832641345 | Lipids / Proteins | C-C stretching (trans-alkane chain) or C-N stretch |
| 3025.64372 | 0.676523296 | Lipids / Proteins | =C-H stretching (unsaturated lipids or aromatics) |
| 1137.67472 | 0.661909992 | Lipids / Proteins | C-C stretching (trans-alkane chain) or C-N stretch |
| 323.861734 | 0.498007892 | Low-frequency skeletal or metal-ligand modes |  |
| 3023.71525 | 0.39057451 | Lipids / Proteins | =C-H stretching (unsaturated lipids or aromatics) |
| 1135.74626 | 0.381787998 | Lipids / Proteins | C-C stretching (trans-alkane chain) or C-N stretch |
| 1633.29069 | 0.380515903 | Proteins | Proteins (β-sheet and disordered protein conformations) |
| 3044.92838 | 0.30309472 | Lipids / Proteins | =C-H stretching (unsaturated lipids or aromatics) |
| 886.974039 | 0.299548841 | Polysaccharides / DNA | C-C stretching or Ribose-phosphate vibrations |
| 1139.60319 | 0.276161019 | Lipids / Proteins | C-C stretching (trans-alkane chain) or C-N stretch |
| 1133.81779 | 0.270030924 | Lipids / Proteins | C-C stretching (trans-alkane chain) or C-N stretch |
| 318.076334 | 0.241626321 | Low-frequency skeletal or metal-ligand modes |  |
| 3179.92106 | 0.238165889 | Proteins / Water | N-H stretching (Amide A) or O-H stretching |
| 867.689371 | 0.227995007 | Polysaccharides / DNA | C-C stretching or Ribose-phosphate vibrations |
| 1542.65275 | 0.208159373 | Proteins (Amide II) | N-H bending and C-N stretching |

**Table 3.** Results from IDH1^WT^ versus IDH1^mut^ classification.

| Raman Frequency | Biochemical Assignment | Molecular Vibration |
| --- | --- | --- |
| ~3345 – 3361 | Proteins / Water | N-H stretching (Amide A) or O-H stretching |
| ~3025 – 3137 | Lipids / Proteins | =C-H stretching (unsaturated lipids/aromatics) |
| ~1617 – 1625 | Proteins (Tyrosine/Phe) | C=C stretching or Tyrosine/Phenylalanine ring mode |
| ~1544 – 1546 | Proteins (Amide II) | N-H bending and C-N stretching |
| ~1216 | Nucleic acids / Proteins | PO<br>asymmetric stretch or Amide III / Tyrosine |
| ~863 – 869 | Polysaccharides / DNA | C-C stretching or Ring breathing modes |
| ~599 | Amino acids / DNA | Phenylalanine ring deformation or C-S stretch |
| ~287 | Lattice / Metal modes | Low-frequency skeletal or metal-ligand vibrations |

**Table 4.** Results from IDH1^mut^ Oligo versus Astro classification.

| Raman Frequency | Biochemical Assignment | Molecular Origin |
| --- | --- | --- |
| ~3349, 3361, 3274 | Proteins / Water | N-H stretching (Amide A) or O-H stretching from hydration |
| ~1413 | Lipids / Proteins | Deformation/bending vibrations. |
| ~1143 | Carbohydrates / Lipids | C-C and C-O stretching (often associated with glucose/glycogen). |
| ~933 - 964 | Proteins / DNA | C-C skeletal stretching (α-helix) or stretching in DNA. |
| ~900 - 906 | Polysaccharides | C-H bending or skeletal vibrations in sugars |
| ~842 | Nucleic acids | C-O-C stretching or Tyrosine (protein) doublet peak |
| ~474 - 505 | Carbohydrates / S-S | Skeletal modes of glycogen or Disulfide (S-S) bonds in proteins |
| ~279 - 312 | Lattice / Metal | Low-frequency skeletal modes or metal-ligand vibrations (e.g., in heme) |

The most prominent differences were observed in the Amide I (1627–1633 cm⁻¹) and Amide II (1542–1544 cm⁻¹) regions, indicating alterations in protein composition and secondary structure. The Amide I band near 1630 cm⁻¹ is frequently associated with β-sheet and disordered protein conformations, suggesting that IDH1 mutation may induce broad changes in the cellular proteome and protein organization.

Several discriminative features were also detected in lipid-associated regions. Peaks at 1120–1139 cm⁻¹ correspond to C–C skeletal vibrations within lipid acyl chains and are commonly associated with membrane lipid order. Additional contributions from the 3023–3139 cm⁻¹ region, assigned to =C–H stretching vibrations of unsaturated lipids, suggest differences in lipid saturation and membrane composition between GBM^mut^ and GBM^WT^ cells. These observations are consistent with reports that mutant IDH1 alters lipid metabolism through the production of the oncometabolite D-2-hydroxyglutarate and subsequent metabolic reprogramming.

Features observed **at** 867–886 cm⁻¹ may reflect contributions from nucleic acids, polysaccharides, or phosphodiester-containing biomolecules, indicating additional changes in cellular biochemical composition. The high-wavenumber feature near 3179 cm⁻¹ likely reflects variations in hydrogen-bonded N–H and O–H groups, suggesting differences in protein hydration or intracellular water organization.

Collectively, these results indicate that the classification model distinguishes GBM^mut^ and GBM^WT^ cells through coordinated changes in protein structure, membrane lipid composition, and cellular metabolic state. The identified Raman features provide evidence that spontaneous Raman spectroscopy captures the biochemical consequences of IDH1 mutation and may serve as a label-free platform for identifying mutation-associated phenotypes in living glioma cells.

We compared these two isogenic cells lines by overlaying the averaged spectra and by also projecting one of the main frequencies onto the whole surface of the cell (Fig. 2).

**Figure 2.**
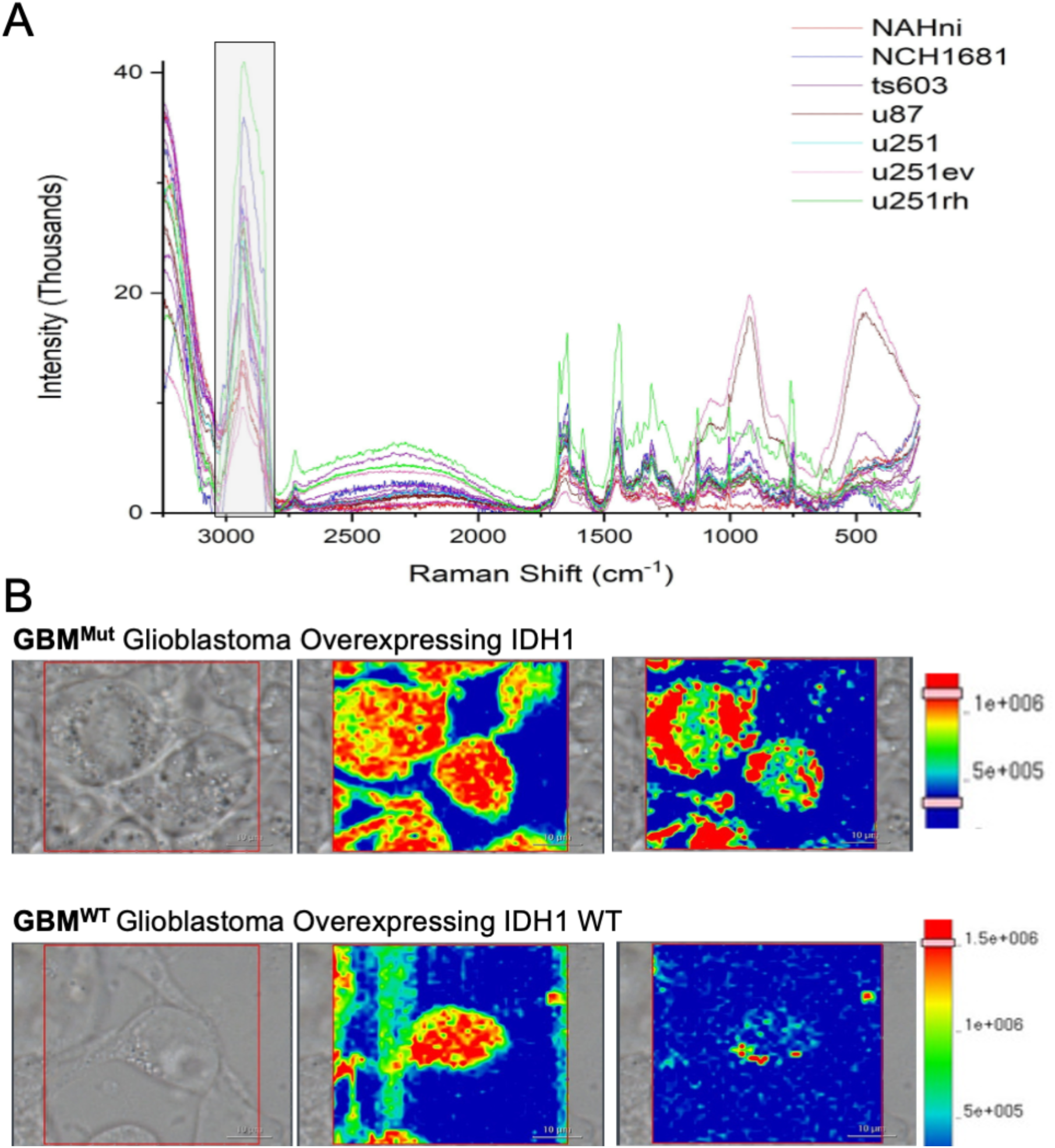
Raman spectra reflect the cellular identity. **A.** Overlayed spectra obtained from cells with different genetic background and different histology. B. Raman spectral maps obtained for two different frequencies for GBM cell line overexpressing IDH1^mut^ or IDH1^WT^ enzymes. Left is shown the optical image for each cell, the middle is the heatmap of one frequency, and the right is the same type of map but for a different frequency.

A second analysis compared patient-derived IDH1 wildtype cells against all IDH1-mutant cells, including oligodendroglioma, astrocytoma, and engineered GBM mutant samples. In this more heterogeneous setting, classification performance decreased but remained consistently above random classification, reaching a balanced accuracy of 0.70. The reduction in performance is expected given the increased biological diversity within the mutant group. The machine learning model identified several Raman frequencies that contributed strongly to the discrimination of IDH1-wildtype and IDH1-mutant glioma cells.

Similar to the GBM^mut^ versus GBM^WT^, the identification of 1544–1546 cm⁻¹ (Amide II) and 1617–1625 cm⁻¹ bands suggest alterations in protein composition and amino acid-associated vibrational modes, consistent with the extensive metabolic and epigenetic reprogramming induced by mutant IDH1. Differences observed in the 3025–3137 cm⁻¹ region, associated with unsaturated lipid vibrations, indicate changes in membrane lipid composition and lipid metabolism, both of which have been linked to IDH1-driven metabolic remodeling^30^. Additional contributions from the 1216 cm⁻¹ band suggest alterations in nucleic acid content or protein secondary structure, while the high-wavenumber features near 3345–3361 cm⁻¹ may reflect changes in protein hydrogen bonding and cellular hydration.

Other discriminatory bands were identified at 3025–3028 cm⁻¹ and 1409 cm⁻¹, corresponding to unsaturated lipid vibrations and CH₂ deformation modes, respectively. These spectral differences indicate alterations in lipid composition and membrane organization, consistent with the profound metabolic remodeling known to occur following acquisition of IDH mutations^22^. Such changes may reflect differences in lipid synthesis, storage, and utilization that accompany the rewiring of central carbon metabolism in IDH-mutant gliomas^48,49,51,53^.

Several protein-associated bands also contributed to classification, including peaks at 867–887 cm⁻¹ and 1213–1217 cm⁻¹, which correspond to protein backbone vibrations and amide III modes. These findings suggest differences in protein composition and secondary structure between IDH-wild-type and IDH-mutant tumors. Furthermore, bands within the high-wavenumber region at 3345–3361 cm⁻¹, associated with O–H and N–H stretching vibrations, indicate differences in hydrogen-bonding interactions and cellular hydration states.

Additionally, in this combined analysis spectral features were bands at 1546 cm⁻¹, 1618–1626 cm⁻¹, and 599 cm⁻¹, which are commonly assigned to heme-containing proteins, particularly cytochrome c. The prominence of these peaks indicates substantial differences in mitochondrial metabolism and cellular redox state between IDH-wild-type and IDH-mutant tumors. Given the central role of cytochrome c in oxidative phosphorylation and apoptotic signaling, these findings suggest that mitochondrial function represents a key biochemical feature distinguishing the two molecular subtypes.

Collectively, these results demonstrate that IDH-wild-type and IDH-mutant gliomas possess distinct Raman spectroscopic fingerprints characterized by alterations in mitochondrial metabolism, lipid organization, protein structure, and cellular microenvironment. The strong contribution of cytochrome c-associated Raman bands suggests that mitochondrial metabolic state is a major determinant of the biochemical differences associated with IDH mutation status and may provide a label-free spectroscopic marker for molecular classification of diffuse gliomas. The identified spectral features provide evidence that Raman spectroscopy captures mutation-associated biochemical phenotypes and may serve as a platform for discovering label-free biomarkers of IDH1 status.

### Raman spectroscopy reveals distinct biochemical signatures within IDH1^mut^ subtypes, further stratification between astrocytoma and oligodendroglioma

To identify biochemical differences between IDH1^mut^ astrocytoma and oligodendroglioma, Raman spectroscopic profiles were analyzed and the spectral features contributing most strongly to tumor classification were identified. Several Raman bands associated with proteins, carbohydrates, lipids, and cellular hydration emerged as major discriminators between the two glioma subtypes. The most influential feature was the Raman band at 906 cm⁻¹, followed by bands at 3349 cm⁻¹ and 308 cm⁻¹. Additional highly ranked features included peaks at 933–964 cm⁻¹, 1143 cm⁻¹, 474 cm⁻¹, 842 cm⁻¹, and 1413 cm⁻¹. The protein-associated region (933–964 cm⁻¹) corresponds to C–C skeletal stretching vibrations characteristic of α-helical protein structures. Additional variations were detected in the Amide A/O–H stretching region (3274–3361 cm⁻¹), suggesting differences in protein hydration state and hydrogen-bonding networks between the two tumor groups.

Bands associated with carbohydrate metabolism were also altered, including peaks at approximately 474–505 cm⁻¹, 900–906 cm⁻¹, and 1143 cm⁻¹, which are commonly assigned to glycogen- and glucose-related vibrational modes. These findings suggest metabolic differences in carbohydrate utilization and storage between astrocytoma and oligodendroglioma.

Differences were further observed in the lipid-associated band at approximately 1413 cm⁻¹, corresponding to CH₂/CH₃ deformation vibrations. Given the lipid-rich nature of oligodendroglial cells, these spectral changes are consistent with distinct lipid metabolic programs between the two glioma subtypes.

Collectively, Raman spectroscopy identified reproducible biochemical differences between astrocytoma and oligodendroglioma, revealing alterations in protein structure, lipid composition, and carbohydrate metabolism that distinguish these tumor entities and may reflect underlying lineage-specific metabolic states.

### Exploratory analysis of normal-cell comparisons

Additional experiments were performed to compare normal cells with oligodendroglioma and astrocytoma cells. These analyses yielded balanced accuracies between 0.75 and 0.81.

Because only six normal-cell samples were available, these results should be interpreted with caution and are reported primarily as exploratory findings. Nevertheless, the observed separation between normal and tumor-derived cells provides further evidence that Raman spectra reflect biologically meaningful differences in cellular composition and state.

## Discussion

The central finding of this study is that live-cell Raman spectroscopy captures two fundamental and biologically distinct determinants of glioma identity: oncogenic mutation status and cellular lineage. Rather than functioning solely as a diagnostic classification tool, Raman spectroscopy revealed biochemical signatures corresponding to the functional consequences of IDH1 mutation and the lineage-specific programs that distinguish oligodendroglioma from astrocytoma. These findings suggest that Raman spectroscopy provides a direct, label-free readout of cellular phenotype in living glioma cells. By combining label-free Raman profiling with computational classification approaches, we identified distinct spectral features that discriminate IDH1-mutant from IDH1-wild-type gliomas as well as oligodendroglioma from astrocytoma. Across the primary classification tasks, balanced accuracies ranged from 0.62 to 0.81 despite substantial biological variability and class imbalance. These findings indicate that Raman spectroscopy captures reproducible biochemical differences associated with cellular phenotype and molecular status.

The strongest performance was observed for the comparison between GBM cells carrying an engineered IDH1 mutation and GBM wildtype cells. This result is particularly interesting because the two groups represent closely related cellular populations that differ primarily in their IDH1 status. The successful discrimination between these groups suggests that Raman spectroscopy can detect biochemical consequences associated with this mutation. Since IDH1 mutations are known to induce substantial metabolic alterations, the observed spectral differences may reflect mutation-associated changes in cellular composition and metabolism.

One of the most striking observations was the prominence of cytochrome c-associated Raman bands among the features distinguishing IDH1-mutant and IDH1-wild-type gliomas. Peaks centered around 1546 cm⁻¹, 1618–1626 cm⁻¹, and 599 cm⁻¹ are commonly associated with heme-containing proteins and mitochondrial electron transport components, suggesting substantial differences in mitochondrial metabolism and cellular redox state between the two groups. These findings are consistent with the extensive metabolic rewiring induced by mutant IDH1. Production of the oncometabolite D-2-hydroxyglutarate alters mitochondrial function, cellular bioenergetics, redox homeostasis, and carbon metabolism, leading to profound biochemical changes that extend beyond the genome and transcriptome^40,41,43,46,49,50,53^. The ability of Raman spectroscopy to detect these differences directly in living cells highlights its potential as a functional readout of tumor metabolism.

The identification of lipid-associated features further supports the existence of distinct metabolic states between IDH1-mutant and IDH1-wild-type tumors. Differences observed in bands corresponding to unsaturated lipid vibrations and CH₂ deformation modes suggest alterations in membrane composition and lipid metabolism. Such findings are consistent with previous studies demonstrating that IDH mutations reshape lipid biosynthesis and membrane remodeling through altered NADPH utilization and central carbon metabolism^40,50^. Importantly, these metabolic consequences are reflected in the biochemical composition of living cells and are therefore accessible to Raman-based interrogation.

The discrimination between astrocytoma and oligodendroglioma cells achieved a balanced accuracy of approximately 0.73. Although lower than the performance obtained for the engineered mutation model, this result demonstrates that Raman spectroscopy captures differences between glioma subtypes arising from distinct developmental lineages.

In contrast to the mitochondrial signatures observed in the IDH1 mutant comparison, classification between lineage-specific IDH1^mut^ glioma subtypes was characterized by differences in protein-associated, carbohydrate-associated, and lipid-associated Raman bands. Multiple discriminatory features were identified within the 933–964 cm⁻¹ region corresponding to protein backbone vibrations, while additional peaks at 474–505 cm⁻¹, 900–906 cm⁻¹, and 1143 cm⁻¹ suggested differences in carbohydrate metabolism and glycogen-associated molecular structures. Together, these findings indicate that astrocytoma and oligodendroglioma possess distinct biochemical programs that extend beyond their shared status as diffuse gliomas.

The lipid-associated signatures identified in the oligodendroglioma–astrocytoma comparison are particularly noteworthy. Although both tumors arise within the diffuse glioma spectrum, oligodendrocytes and astrocytes perform fundamentally different physiological functions within the central nervous system. Oligodendrocytes are specialized for myelin production and lipid synthesis, whereas astrocytes serve broader metabolic and homeostatic roles. These developmental programs persist, at least in part, following malignant transformation and contribute to differences in cellular composition, energy utilization, and biosynthetic activity. The ability of Raman spectroscopy to detect these differences suggests that lineage identity leaves a measurable biochemical imprint that remains detectable in living tumor cells, in addition to IDH1^mut^-associated metabolic reprogramming.

Another notable finding was the contribution of high-wavenumber bands associated with O–H and N–H stretching vibrations. These features likely reflect differences in cellular hydration, hydrogen bonding, and macromolecular organization. Such parameters are difficult to assess using conventional molecular techniques but can influence protein conformation, organelle function, and cellular metabolism. Their identification as discriminatory features further illustrate the breadth of biological information contained within Raman spectra.

Methodologically, an important aspect of the present workflow is the use of strict sample-level validation. Although the dataset contains nearly 700,000 spectra, the effective number of independent observations is determined by the number of biological samples rather than the number of spectra. To avoid information leakage, all spectra originating from a given sample were assigned to the same cross-validation fold and final predictions were generated using sample-level majority voting. This evaluation protocol provides a more realistic assessment of model performance than spectrum-level validation.

Another important component of the workflow is the use of RADAR preprocessing^27,28^. Conventional Raman preprocessing pipelines typically rely on a sequence of independently designed procedures for baseline correction, denoising, and artifact removal. In contrast, RADAR performs simultaneous decomposition of each spectrum into baseline, cosmic-ray, noise, and peak components using deep neural networks trained on synthetic Raman data. By retaining only the reconstructed peak component, the method removes several sources of spectral variability in a single processing step while preserving the biochemical information relevant for classification. The use of a consistent preprocessing framework across all experiments also reduces variability introduced by manual selection of preprocessing parameters.

One limitation is that the current study focuses on a limited number of cell lines acquired under controlled experimental conditions. While this setting enables investigation of specific biological questions, additional validation will be required to determine the extent to which the identified spectral signatures generalize to independent experiments and acquisition sessions. Another limitation is that individual Raman bands can arise from multiple molecular contributors, and therefore biochemical assignments should be interpreted in the context of broader spectral patterns rather than as direct measurements of individual molecules.

Therefore, future work will focus on few directions. The first is a detailed feature-importance analysis of the trained XGBoost models to identify the spectral regions responsible for the observed classifications. The second is validation using an independent set of Raman measurements acquired separately from the data used for model development. In addition, while the present study identifies robust spectral signatures associated with mutation status and lineage identity, future integration with metabolomic, lipidomic, and transcriptomic datasets will be important for defining the precise molecular pathways responsible for these signatures. Together, these analyses will provide a deeper understanding of the biological basis of the observed classifications and further evaluate the robustness of the proposed workflow.

Taken together, our findings support a model in which IDH mutation and lineage identity represent two orthogonal biological axes that shape glioma phenotype. IDH1 mutation primarily influences cellular metabolism, mitochondrial function, and redox homeostasis, whereas lineage identity determines broader aspects of cellular architecture, protein composition, lipid organization, and metabolic specialization. Remarkably, Raman spectroscopy was able to resolve both axes simultaneously using only intrinsic biochemical information. This capability distinguishes Raman spectroscopy from conventional molecular assays, which typically interrogate either genotype or histology independently.

The use of live-cell Raman spectroscopy provides several important advantages. Because measurements are non-destructive and do not require fixation, staining, or exogenous probes, cellular physiology remains largely unperturbed during analysis. Consequently, Raman spectroscopy enables direct characterization of living cells while preserving the dynamic biochemical states that are often lost during conventional sample processing. This feature may be particularly valuable for studying metabolic plasticity, treatment response, co-culture systems, patient-derived organoid models, and state transitions in glioma cells.

## Acknowledgements

The authors would like to thank Alan and Ashley Dabbiere for their generous donation which enabled the acquisition of the Leica Stellaris 8 CRS instrument. This research was supported by the National Institutes of Health Intramural Research Program through an NCI FLEX award to A.L. and M.L. entitled “Live cell metabolism via Raman imaging microscopy.”

## Funding

This research was supported by the Intramural Research Program of the National Institutes of Health (NIH): ZIA BC011711-09 “Develop RAMAN imaging to visualize metabolism in cell lines and tissue”. The contributions of the NIH author(s) are considered Works of the United States Government. This project has been funded in whole or in part with Federal funds from the National Cancer Institute, National Institutes of Health, Department of Health and Human Services. The content of this publication does not necessarily reflect the views or policies of the Department of Health and Human Services, nor does mention of trade names, commercial products, or organizations imply endorsement by the U.S. Government

## Notes

### Competing Interest Statement

The authors have declared no competing interest.

